# Reorganizing Niche Architecture Still Preserves Organ Function in the Hair Follicle

**DOI:** 10.1101/2022.09.06.506764

**Authors:** Haoyang Wei, Shuangshuang Du, Jeeun Parksong, H. Amalia Pasolli, Sergi Regot, Lauren E. Gonzalez, Tianchi Xin, Valentina Greco

## Abstract

Stem cells’ ability to build and replenish tissues depends on support from their niche. While niche architecture varies across different organs, the functional importance of niche architecture is unclear. During hair follicle growth, multipotent epithelial progenitors build hair via crosstalk with their remodeling fibroblast niche, the dermal papilla, providing a powerful model to functionally interrogate different niche architectures. Through intravital imaging, we show that dermal papilla fibroblasts remodel both individually and collectively to form a polarized, structurally robust niche. Polarized TGFβ signaling precedes structural niche polarity, and loss of TGFβ signaling in dermal papilla fibroblasts leads them to progressively lose their stereotypic architecture and instead surround the epithelium. The reorganized niche relocates multipotent progenitors, but nevertheless supports their proliferation and differentiation. However, progenitor differentiation is completed prematurely, resulting in compromised hair production. Overall, our results reveal that niche architecture optimizes organ efficiency, but is not absolutely essential for organ function.

## Introduction

Interaction between stem cells and their niche lays the foundation for organ architecture and function (Xin et al., 2016, Fuchs and Blau, 2020). The niche creates a microenvironment with the correct signaling and physical characteristics to support stem cell survival and production of differentiated daughter cells (O’Brien and Bilder, 2013, Lane et al., 2014). Niches across mammalian tissues take a variety of architecture types, including ensheathing stem cells (intestine, lung, mammary gland, etc) and enclosed by stem cells (hair follicle, tooth, tongue, etc) (Plikus et al., 2021, McCarthy et al., 2020, Goodwin and Nelson, 2020, Mao and Prockop, 2012, Mistretta and Bradley, 2021). This implies that niche architecture is important for stem cell function, but few tools exist to test the function of niche architecture *in vivo*.

The hair follicle is a useful model for studying the impact of organ architecture on function because hair follicles recurrently grow and regress in each hair cycle throughout adulthood (MuÈller-RoÈver et al., 2001). This remarkable cycling is achieved by concurrent rearrangement of hair follicle stem cells and associated niche fibroblasts in the dermal papilla and dermal sheath (Fig.S1A) (Chi et al., 2013, Morgan, 2014, Rahmani et al., 2014, Heitman et al., 2020). Dermal papilla fibroblasts form a single cluster at the bottom of the hair follicle and remodel dramatically during hair follicle growth (Driskell et al., 2011). The dermal papilla is first enclosed by the proliferating progeny of hair stem cells, a pool of multipotent progenitors (also called hair matrix). The dermal papilla then significantly elongates with negligible fibroblast proliferation while secreting growth factors to instruct the proliferation and differentiation of multipotent progenitors (Tobin et al., 2003, Morgan, 2014). Consequently, seven epithelial lineages arranged in a concentric organization are generated, fueling hair production (Fig.1A). Thus, dermal papilla remodeling during hair follicle regeneration provides an outstanding model to study how different niche architectures impact their crosstalk with stem cells and overall organ function.

**Figure 1.**
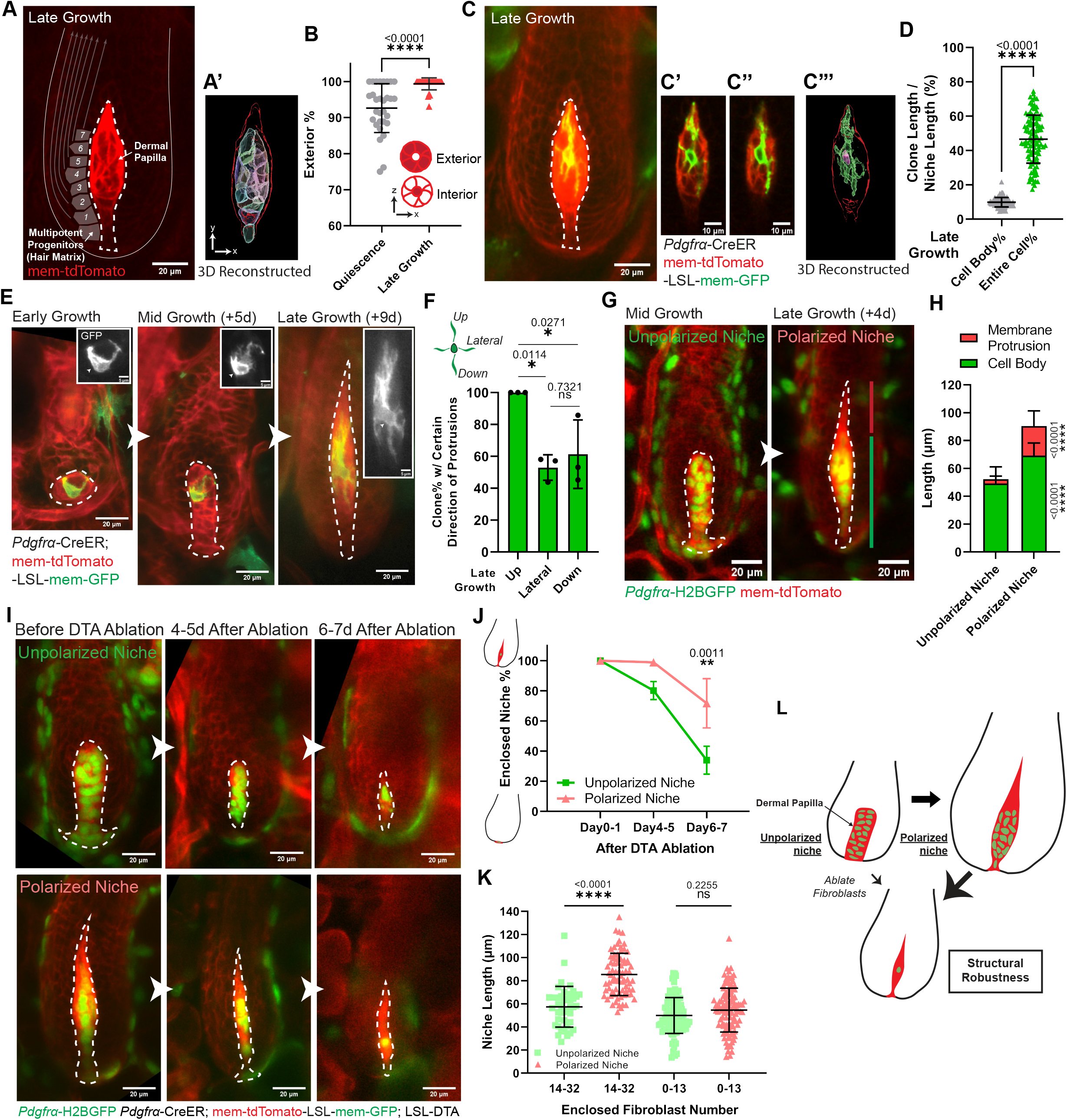
Niche cells undergo collective polarization to confer a structurally robust architecture. A. Intravitally imaged dermal papilla at the late-growth stage of hair follicles (n=3 mice). Dermal papilla fibroblasts are labeled with membrane-tdTomato, and the entire dermal papilla is outlined. Gray squares illustrate adjacent multipotent progenitors, with arrows indicating their differentiation routes. (A’) An entire dermal papilla is 3D reconstructed based on its membrane-tdTomato signal. Within the red outlined dermal papilla, cell bodies of individual fibroblasts are rendered into surfaces with different colors. B. Percentage of fibroblasts whose cell bodies are located at the exterior of the dermal papilla. Exterior-localized is defined as any fibroblasts whose cell body directly contacts the dermal papilla edge; interior-localized is defined as a fibroblast cell body surrounded by cell bodies of other fibroblasts. n=30 quiescent and 30 late-growth hair follicles from 3 mice. C. Intravital imaging of a single fibroblast at late-growth stage (n=3 mice). (C’-C’’) Long membrane protrusions extend through the cell bodies of other fibroblasts. (C’’’) The single fibroblast is 3D reconstructed based on its membrane-GFP signal. Within the red-outlined dermal papilla, the cell body of this fibroblast is rendered into a pink surface and its entire membrane in a green surface. D. Coverage percentage of dermal papilla fibroblast clones, as the length of fibroblast cell body or entire membrane relative to length of entire dermal papilla at late-growth. n=103 dermal papilla fibroblast clones at late-growth from 3 mice. E. One representative dermal papilla fibroblast longitudinally imaged from early to late-growth (n=3 mice). Insets highlight membrane-GFP (gray) and fibroblast cell body (arrowheads). F. Percentage of dermal papilla fibroblasts harboring certain direction of membrane protrusions (upward, lateral, downward) at late-growth. n=77 dermal papilla fibroblast clones at late-growth from 3 mice. G. The same dermal papilla longitudinally imaged from mid to late-growth. The membrane protrusion compartment (red line) on top of the cell body compartment (green line) organizes an architecturally polarized niche at late-growth, in contrast to an unpolarized niche at mid-growth. H. Length of different dermal papilla compartments at mid-growth (Late anagen 2-3b, unpolarized niche) and late-growth stage (Anagen 3c-6, polarized niche). n=117 unpolarized dermal papillae and 102 polarized dermal papillae from 3 mice. I. The same unpolarized and polarized dermal papillae longitudinally imaged before and after diphtheria toxin (DTA) induced fibroblast ablation. Ablation is achieved by tamoxifen activated Cre in fibroblasts (*Pdgfrα*-CreER) and LSL-DTA. J. Percentage of unpolarized and polarized dermal papillae that remain enclosed after fibroblast ablation. For unpolarized dermal papillae, n=3 mice (68 hair follicles at Day0-1, 68 hair follicles at Day4-5, and 66 hair follicles at Day6-7); for polarized dermal papillae, n=3 mice (102 hair follicles at Day0-1, 108 hair follicles at Day4-5, and 89 hair follicles at Day6-7). K. Length of entire dermal papillae that remain enclosed after ablation. Based on the reported critical threshold of dermal papilla fibroblast number (Chi et al., 2013), we classified dermal papillae as either with abundant (14 to 32) or few (0 to 13) enclosed fibroblasts. For unpolarized dermal papillae, n=42 hair follicles (14-32 fibroblasts) and 99 hair follicles (0-13 fibroblasts) from 3 mice; For polarized dermal papillae, n=77 hair follicles (14-32 fibroblasts) and 110 hair follicles (0-13 fibroblasts) from 3 mice. L. Schematic of forming a polarized dermal papilla niche architecture that is structurally robust even after fibroblast ablation. Green indicates dermal papilla fibroblast nuclei, and red indicates their membranes. All representative images have scale bar sizes indicated. Images are single optical z slice in Fig.1A, C’-C’’, and summed optical z slices in Fig.1C, E, G, I. Dermal papillae are dash-lined. In Fig.1C, E, single fibroblast is labeled by mosaic recombined membrane-GFP (mTmG) under the fibroblast driven CreER (*Pdgfrα*-CreER). In Fig.1G, I, fibroblast nuclei are labeled in green by *Pdgfrα*-H2BGFP, and cell membranes are in red by membrane-tdTomato. All data are presented as mean ± S.D. Unpaired two-tailed t-test is used in Fig.1B, D, H. Tukey’s multiple comparisons test is used in Fig.1F, J, K. See also Figure S1-2, Supplemental Item 1.

As multipotent progenitors reposition along the dermal papilla, they flexibly adopt different fates depending on the timing and location of their disassociation from the niche (Legue and Nicolas, 2005, Legue et al., 2010, Mesler et al., 2017, Xin et al., 2018). Signaling crosstalk between the dermal papilla and multipotent progenitors has been relatively well characterized (Rendl et al., 2005, Greco et al., 2009, Sennett and Rendl, 2012, Hsu et al., 2014, Rezza et al., 2016, Yang et al., 2017, Adam et al., 2018, Harshuk-Shabso et al., 2020), but the direct impacts of the dermal papilla’s three-dimensional (3D) organization on hair follicle growth is not well understood. To address this question, we leveraged our ability to manipulate and longitudinally track the same hair follicle niche fibroblasts in live mice (Fig.S1B) (Pineda et al., 2015). We found that during hair follicle growth, dermal papilla fibroblast membranes remodel to form a polarized architecture that remains structurally robust even in the face of fibroblast death. Just before architectural polarization, TGFβ signaling within the dermal papilla is activated in a polarized pattern, and fibroblast TGFβ signaling is required for retention of the dermal papilla’s enclosed architecture. Finally, by reorganizing the niche through blocking TGFβ signaling, we discovered that the enclosed architecture of the dermal papilla dictates the location of multipotent progenitors for efficient differentiation and correct hair production. However, the reorganized niche nevertheless still preserves organ function by supporting the basic proliferation and differentiation of progenitors. Taken together, we uncovered how the structural features of a remodeling niche are formed and optimize stem cell activities and organ function.

## Results

### Niche cells undergo collective polarization to confer a structurally robust architecture

The dermal papilla is a small cluster when it supports epithelial proliferation at early growth stages, and elongates as it begins supporting epithelial differentiation at later growth stages (Fig.S1A) (MuÈller-RoÈver et al., 2001). To understand whether fibroblasts also change their organization within the dermal papilla during remodeling, we labeled them with membrane-tdTomato and 3D reconstructed their cell bodies (Fig.1A-A’). At the quiescent stage, we found that the dermal papilla contains a population of interior-localized fibroblasts surrounded by exterior-localized fibroblasts (Fig.1B). In contrast, at the late growth stage, nearly all fibroblasts position their cell bodies along the exterior of the dermal papilla, directly interfacing with adjacent multipotent progenitors (Fig.1A-B; SI.1-Movie). This suggests that the dermal papilla at late growth is organized to optimally communicate with multipotent progenitors.

To understand how fibroblast morphology contributes to these organizational changes within the dermal papilla, we resolved individual fibroblasts by mosaic labeling with membrane-GFP (*Pdgfrα*-CreER; mTmG). Surprisingly, we found that at the late growth stage, individual dermal papilla fibroblasts extend long membrane protrusions across the entire niche (Fig.1C-D; SI.1-Movie), allowing them to contact distant fibroblasts and multipotent progenitors. This contrasts with the much shorter membrane protrusions on dermal papilla fibroblasts at mid-growth (Fig.1E; Fig.S2A-B). Interestingly, at the late growth stage, only a subset have lateral or downward-oriented membrane protrusions but all fibroblasts have upward-oriented protrusions (Fig.1F; Fig.S2C). By differentially labeling all fibroblast nuclei and cell membranes (*Pdgfrα*-H2BGFP; mem-tdTomato), we found that dermal papilla fibroblast membrane protrusions are close to their cell bodies at mid-growth, but at the late growth stage, the dermal papilla develops a distinct membrane protrusion compartment over its cell body compartment (Fig.1G-H). Altogether, this indicates that dermal papilla fibroblasts collectively form an upward polarized niche architecture at the late growth stage by extending membrane protrusions and spatially compartmentalizing them from their cell bodies.

To better understand the structural properties of dermal papilla architecture throughout its polarization process, we asked how different niche architectures respond to fibroblast removal. We genetically induced fibroblast apoptosis by expressing diphtheria toxin (*Pdgfrα*-H2BGFP; *Pdgfrα*-CreER; mTmG; LSL-DTA) at the unpolarized (mid-growth) and polarized (late-growth) niche stages, and revisited the same dermal papillae over time. The loss of *Pdgfrα*-H2BGFP positive nuclei confirmed fibroblast death at similar kinetics for both niche stages (Fig.1I; Fig.S2D-E). Surprisingly, we found that the majority of polarized niches remained enclosed by the epithelium six to seven days after fibroblast ablation, in contrast to the earlier and more frequent loss of enclosed architecture in unpolarized niches after fibroblast ablation (Fig.1J). Notably, in both polarized and unpolarized dermal papillae that remain enclosed despite losing more than half of their fibroblasts, fibroblast membranes are still extended to occupy a similar overall dermal papilla length (Fig.1I, 1K). These data together suggest that niche architecture becomes more structurally robust as it polarizes at the single-cell and tissue levels during hair follicle growth (Fig.1L).

### Niche architecture and location are actively maintained through fibroblast TGFβ signaling

The differential effects of fibroblast ablation on unpolarized and polarized niches suggest a link between niche architectural polarity and maintenance. The expression of TGFβ receptor 1 (TGFβRI) begins in dermal papilla fibroblasts right before architectural polarization (Paus et al., 1997), making it a promising candidate mediator of dermal papilla polarity and/or structural robustness. To investigate the role of TGFβ signaling in the dermal papilla, we first examined when and where TGFβ signaling is activated in dermal papilla fibroblasts. We built a TGFβ reporter mouse line by fluorescently tagging endogenous Smad4 (mNeonGreen-Smad4, Fig.S3A-B), which has been reported to accumulate in the nucleus upon TGFβ signaling (Warmflash et al., 2012). Mouse embryonic fibroblasts from our reporter mouse showed nuclear localization of mNeonGreen-Smad4 in response to the addition of TGFβ (Fig.S3C), confirming that this mouse line reflects TGFβ signaling. Excitingly, as we tracked signaling activity *in vivo*, we found that nuclear Smad4 is first detectable in upper dermal papilla fibroblasts in the unpolarized niche, and this signaling polarization is sustained as the dermal papilla becomes structurally polarized (Fig.2A).

**Figure 2.**
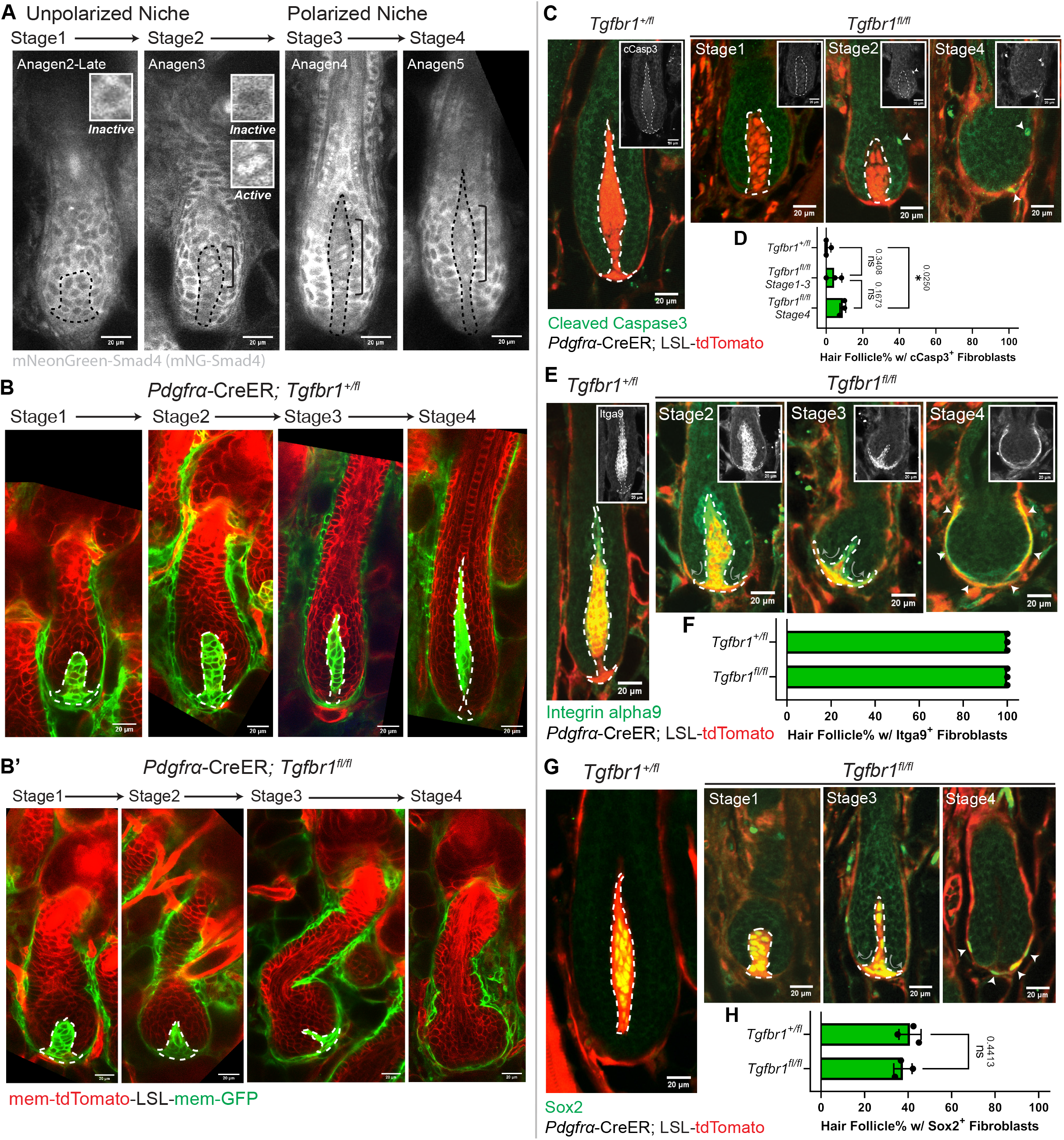
Niche architecture and location are actively maintained through fibroblast TGFβ signaling. A. Intravital imaging of our TGFβ signaling reporter, mNeonGreen-Smad4 (gray), in hair follicles at different stages (n=3 mice). Nuclear-localized Smad4 are present in dermal papilla fibroblasts within the bracket. Insets show inactive (cytoplasmic) or active (nuclear) Smad4 signals. B. Intravital imaging of *Tgfbr1*^*+/fl*^ or (B’) *Tgfbr1*^*fl/fl*^ hair follicles from mid to late-growth (Stage1-4) after Cre induction at the quiescent stage (n=3 mice). Fibroblast membranes are in green membrane-GFP and other membranes are in red membrane-tdTomato. C. Immunostaining for cleaved Caspase3 (cCasp3, green) in thick back skin sections to detect apoptosis at different stages. Arrowheads indicate cCasp3 signal. Insets show cCasp3 in gray. D. Percentage of *Tgfbr1*^*+/fl*^ and *Tgfbr1*^*fl/fl*^ hair follicles at different stages containing cCasp3^+^ fibroblasts. n=3 *Tgfbr1*^*+/fl*^ mice (280 hair follicles at Stage4), n=3 *Tgfbr1*^*fl/fl*^ mice (49 hair follicles at Stage1-3, 81 hair follicles at Stage4). Tukey’s multiple comparisons test. E. Immunostaining for Integrin alpha9 (Itga9, green) in thick back skin sections to detect dermal papilla fibroblast identity at different stages. Arrowheads indicate Itga9^+^ regions in Stage4. Arrows indicate the reorganizing direction of dermal papilla fibroblasts. Insets show Itga9 in gray. F. Percentage of *Tgfbr1*^*+/fl*^ and *Tgfbr1*^*fl/fl*^ hair follicles containing Itga9^+^ fibroblasts. n=3 *Tgfbr1*^*+/fl*^ mice (234 late-growth hair follicles), n=3 *Tgfbr1*^*fl/fl*^ mice (144 hair follicles from stages shown). G. Immunostaining for Sox2 (green) in thick back skin sections to detect dermal papilla fibroblast identity at different stages. Arrowheads indicate Sox2^+^ regions in Stage4. Arrows indicate the reorganizing direction of dermal papilla fibroblasts. H. Percentage of *Tgfbr1*^*+/fl*^ and *Tgfbr1*^*fl/fl*^ hair follicles containing Sox2^+^ fibroblasts. n=3 *Tgfbr1*^*+/fl*^ mice (594 late-growth hair follicles), n=3 *Tgfbr1*^*fl/fl*^ mice (226 hair follicles from stages shown). Unpaired two-tailed t-test. All representative images are single optical z slice with scale bar sizes indicated. Dermal papillae are dash-lined in all images. In Fig.1C, E, G, fibroblasts are labeled in red by *Pdgfrα*-CreER; LSL-tdTomato for both *Tgfbr1*^*+/fl*^ and *Tgfbr1*^*fl/fl*^. All data are presented as mean ± S.D. See also Figure S3-4.

The spatiotemporal symmetry between polarized signaling activity and polarized structural architecture prompted us to functionally examine the consequences of TGFβ signaling loss on dermal papilla architecture. To this end, we knocked out TGFβRI in fibroblasts from the quiescent stage via Cre induction at postnatal day 18-20 and used fluorescent reporters to identify fibroblasts that expressed Cre (*Pdgfrα*-CreER; *Tgfbr1*^*fl/fl*^ combined with either membrane-GFP or cytoplasmic tdTomato reporter). Revisits of the same mice showed that *Tgfbr1*^*fl/fl*^ mutant niches proceeded to the unpolarized stage like *Tgfbr1*^*+/fl*^ controls (Fig.2B-B’, Stage1; Fig.S4A). However, they failed both to elongate and to polarize, and gradually lost their enclosed architecture (Fig.2B-B’, Stage2-4). Negligible fibroblast apoptosis was detected prior to loss of tdTomato^+^ (*Tgfbr1*^*fl/fl*^) fibroblast enclosure (Fig.2C-D, Stage1-2), indicating that fibroblast apoptosis is not the cause of niche architecture loss.

To better understand the effects of TGFβRI deletion prior to loss of dermal papilla architecture, we asked whether dermal papilla fibroblasts lose their identity markers. We stained for two dermal papilla identity markers: integrin alpha9 and Sox2 (Driskell et al., 2009, Tsai et al., 2010). Surprisingly, we found that dermal papilla fibroblasts retained identity markers in *Tgfbr1*^*fl/fl*^ mutants. In particular, the position of integrin alpha9 and Sox2 positive fibroblasts progressively transitioned from enclosed within the epithelium to ensheathing around the epithelium (Fig.2E-H). This niche reorganization indicates that dermal papilla niche architecture and location are actively maintained, and this maintenance depends on fibroblast TGFβ signaling.

### A reorganized niche relocates multipotent progenitors but largely supports their function

The reorganized niche architecture in the *Tgfbr1*^*fl/fl*^ mutant provided us with an excellent opportunity to investigate how epithelial multipotent progenitors are impacted by niche architecture, which has been understudied compared to their signaling crosstalk. We first interrogated the proliferation status of multipotent progenitors in the *Tgfbr1*^*fl/fl*^ mutant by staining for the proliferation marker Ki67. Despite the changed niche architecture, mutant hair follicles had an overall bulb-like structure containing a pool of proliferative epithelial cells, similar to control hair follicles (Fig.3A-B). However, only epithelial cells in the outer bulb layers were proliferative in mutant hair follicles, indicating a spatial reorganization of the bulb region (Fig.3C). Furthermore, the pattern of cells in the proliferation zone expressing Lef1, a multipotent progenitor marker (Mesler et al., 2017), also redistributed to the outer epithelial layers upon niche relocation (Fig.3D-E), further demonstrating that multipotent progenitors remain proliferative, but their location depends on niche organization.

**Figure 3.**
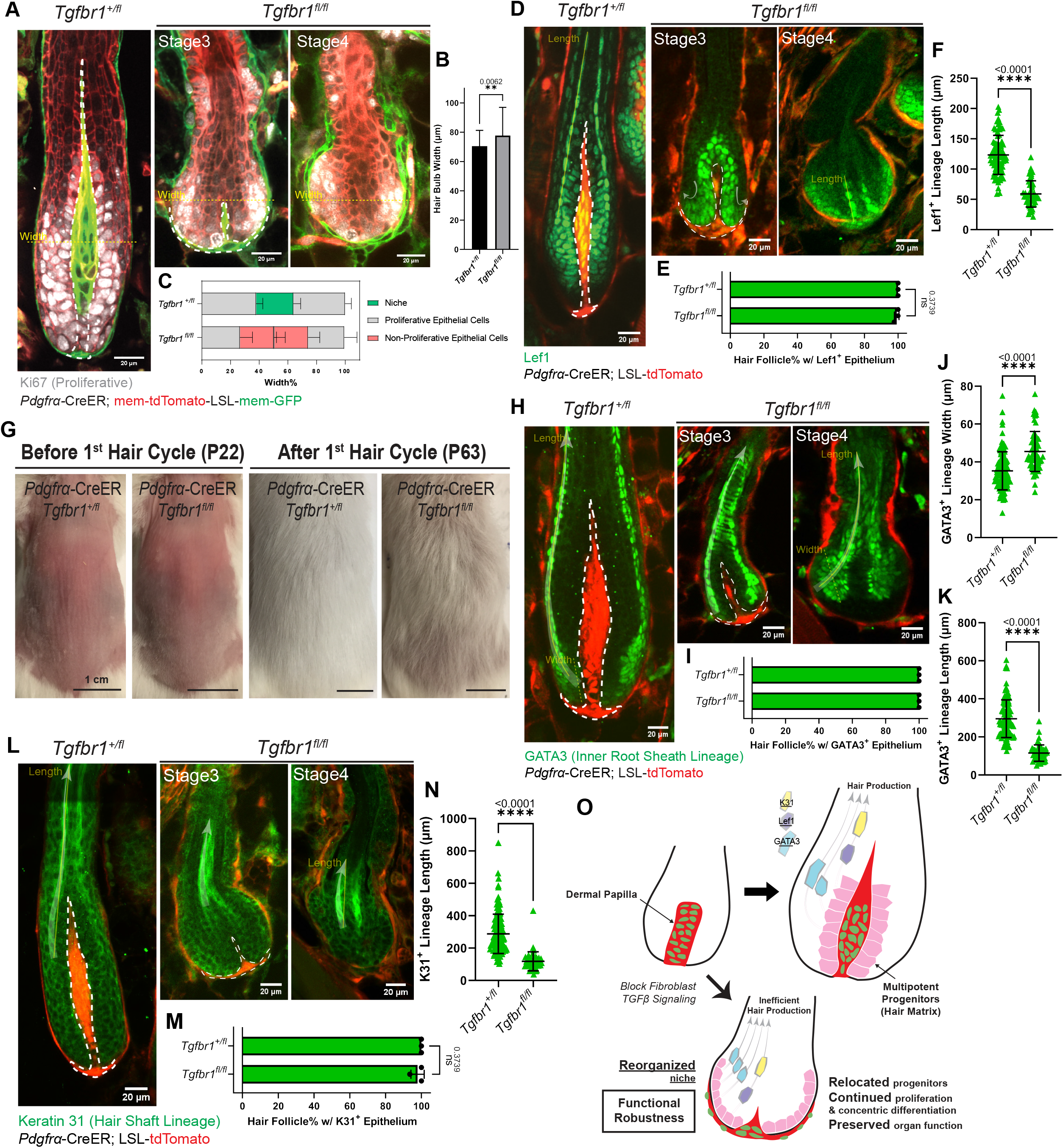
A reorganized niche relocates multipotent progenitors but largely supports their function. A. Immunostaining for Ki67 (gray) in ear skin whole mounts to detect proliferative epithelial cells in hair follicles at different stages. Fibroblast membranes are in green (*Pdgfrα*-CreER; membrane-GFP) and other membranes are in red (membrane-tdTomato). B. Hair bulb maximal width in *Tgfbr1*^*+/fl*^ and *Tgfbr1*^*fl/fl*^ at the location of yellow dash-lines in Fig.3A. n=75 hair follicles from three *Tgfbr1*^*+/fl*^ mice, n=62 hair follicles from three *Tgfbr1*^*fl/fl*^ mice. C. Spatial distribution of different cell populations (niche, proliferative epithelial cells, non-proliferative epithelial cells) in the bulb region, measured as the width percentage of each population at the location of maximal hair bulb width (yellow dash-lines in Fig.3A). n=75 hair follicles from three *Tgfbr1*^*+/fl*^ mice, n=62 hair follicles from three *Tgfbr1*^*fl/fl*^ mice. D. Immunostaining for Lef1 (green) in thick back skin sections to detect multipotent progenitors and Wnt/β-catenin signaling at different stages. The relocation directions are marked by arrows in light gray. E. Percentage of hair follicles in *Tgfbr1*^*+/fl*^ and *Tgfbr1*^*fl/fl*^ mice containing Lef1^+^ epithelial populations in the bulb region. n=3 *Tgfbr1*^*+/fl*^ mice (215 late-growth hair follicles), n=3 *Tgfbr1*^*fl/fl*^ mice (93 hair follicles from stages shown). F. Length of Lef1^+^ differentiating lineage in Stage4 of *Tgfbr1*^*+/fl*^ and *Tgfbr1*^*fl/fl*^ hair follicles along the differentiation routes (yellow solid line in Fig.3D). n=81 hair follicles from three *Tgfbr1*^*+/fl*^ mice, n=49 hair follicles from three *Tgfbr1*^*fl/fl*^ mice. G. Representative photos of *Tgfbr1*^*+/fl*^ and *Tgfbr1*^*fl/fl*^ mice with back skin shaved at the quiescent stage, and revisited after an entire first hair cycle. H. Immunostaining for GATA3 (green) in thick back skin sections to detect the differentiation of inner root sheath lineages (Cuticle and Huxley’s layer) at different stages. The differentiation routes are marked with arrows in light gray. I. Percentage of hair follicles in *Tgfbr1*^*+/fl*^ and *Tgfbr1*^*fl/fl*^ mice containing GATA3^+^ epithelial populations in concentric organizations. n=3 *Tgfbr1*^*+/fl*^ mice (328 late-growth hair follicles), n=3 *Tgfbr1*^*fl/fl*^ mice (103 hair follicles from stages shown). J. Width of GATA3^+^ differentiating lineage in Stage4 of *Tgfbr1*^*+/fl*^ and *Tgfbr1*^*fl/fl*^ at the beginning of differentiation routes (yellow dash line in Fig.3H). n=109 hair follicles from three *Tgfbr1*^*+/fl*^ mice, n=59 hair follicles from four *Tgfbr1*^*fl/fl*^ mice. K. Length of GATA3^+^ differentiating lineage in *Tgfbr1*^*+/fl*^ and *Tgfbr1*^*fl/fl*^ Stage4 hair follicles along the differentiation routes (yellow solid line in Fig.3H). n=112 hair follicles from three *Tgfbr1*^*+/fl*^ mice, n=51 hair follicles from four *Tgfbr1*^*fl/fl*^ mice. L. Immunostaining for Keratin 31 (K31, green) in thick back skin sections to detect the differentiation of hair shaft lineage (Cortex) at different stages. The differentiation routes are marked with arrows in light gray. M. Percentage of hair follicles in *Tgfbr1*^*+/fl*^ and *Tgfbr1*^*fl/fl*^ mice containing K31^+^ epithelial populations in concentric organizations. n=3 *Tgfbr1*^*+/fl*^ mice (243 late-growth hair follicles), n=3 *Tgfbr1*^*fl/fl*^ mice (109 hair follicles from stages shown). N. Length of K31^+^ differentiating lineage in the Stage4 of *Tgfbr1*^*+/fl*^ and *Tgfbr1*^*fl/fl*^ along the differentiation routes (yellow solid line in Fig.3L). n=132 hair follicles from three *Tgfbr1*^*+/fl*^ mice, n=44 hair follicles from three *Tgfbr1*^*fl/fl*^ mice. O. Schematic illustrating how reorganized niche architecture continues supporting relocated proliferative multipotent progenitors. However, progenitor differentiation is rerouted and completes prematurely for less efficient hair production. All representative images are single optical z slice with scale bar sizes indicated. Dermal papillae are dash-lined in all images. In Fig.1D, H, L, fibroblasts are labeled in red by *Pdgfrα*-CreER; LSL-tdTomato for both *Tgfbr1*^*+/fl*^ and *Tgfbr1*^*fl/fl*^. All data are presented as mean ± S.D. and analyzed with unpaired two-tailed t-test. See also Figure S4.

We next asked whether the re-organized niche and epithelial progenitors can still support hair production. We shaved the mouse back skin at the quiescent stage (postnatal day 18-20) and tracked hair regrowth after an entire hair cycle. Interestingly, we found that hair shafts were still produced in *Tgfbr1*^*fl/fl*^ mutant mice, though with less coverage than *Tgfbr1*^*+/fl*^ controls (Fig.3G; Fig.S4B). To understand what caused this decreased hair coverage, we stained for markers of two differentiation lineages, inner root sheath (GATA3) and hair shaft (Keratin 31) (Yang et al., 2017). Notably, both GATA3^+^ and Keratin 31^+^ lineages were still present and remained in a concentric organization like in *Tgfbr1*^*+/fl*^ controls (Fig.3H-I; Fig.3L-M). However, GATA3^+^ epithelial cells in the *Tgfbr1*^*fl/fl*^ mutants expanded their differentiation pool (indicated by increased lineage width, Fig.3J) and adopted a concave path rather than a convex path (Fig.3H, arrow). This indicates that multipotent progenitors spatially reroute their differentiation path upon niche reorganization. Furthermore, both GATA3^+^ and Keratin 31^+^ lineages showed precocious loss of their differentiation markers along the differentiation path (indicated by decreased lineage length, Fig.3H, 3L, Stage4; Fig.3K, 3N). Similarly, Lef1 activity above the proliferation zone, which reflects Wnt signaling-induced differentiation (Adam et al., 2018, Choi et al., 2013), was precociously lost in *Tgfbr1*^*fl/fl*^ mutant differentiating progenitors (Fig.3D, Stage4; Fig.3F). These results together indicate that multipotent progenitors under the instruction of a reorganized niche proliferate, initiate differentiation correctly but fail to maintain their differentiation program and produce hair shafts less efficiently.

Overall, we elucidated that dermal papilla niche architecture dictates multipotent progenitor location and differentiation for efficient hair production, which is possible but compromised in a sheath-like niche architecture (Fig.3O).

## Discussion

The niche constructs a specific microenvironment of signaling and mechanical cues that support stem cell survival and function. While the transcriptional signatures and signaling activities of niche cells have received considerable attention, the functional importance of niche 3D architecture remains elusive across a number of organs. The hair follicle niche, dermal papilla, is an ideal model to study how niche architecture impacts stem cell organization and function because it undergoes remarkable architectural remodeling during hair growth. Here, by longitudinal intravital imaging of individual fibroblasts and the entire niche architecture, we showed that dermal papilla fibroblasts collectively remodel cell body positions and membrane protrusions to form an upward polarized and structurally robust niche architecture. TGFβ signaling is required for retention of its enclosed architecture. Most surprisingly, we found that a reorganized dermal papilla can still sustain hair progenitor proliferation and concentric organization of differentiation. However, hair progenitor differentiation is completed prematurely and hair shaft formation is compromised (Fig.4). Overall, our discovery that the hair follicle niche progressively acquires structural and functional robustness, such that it can continue supporting progenitor function even when re-organized, provides insight into general regulatory and functional principles of niche architecture.

**Figure 4.**
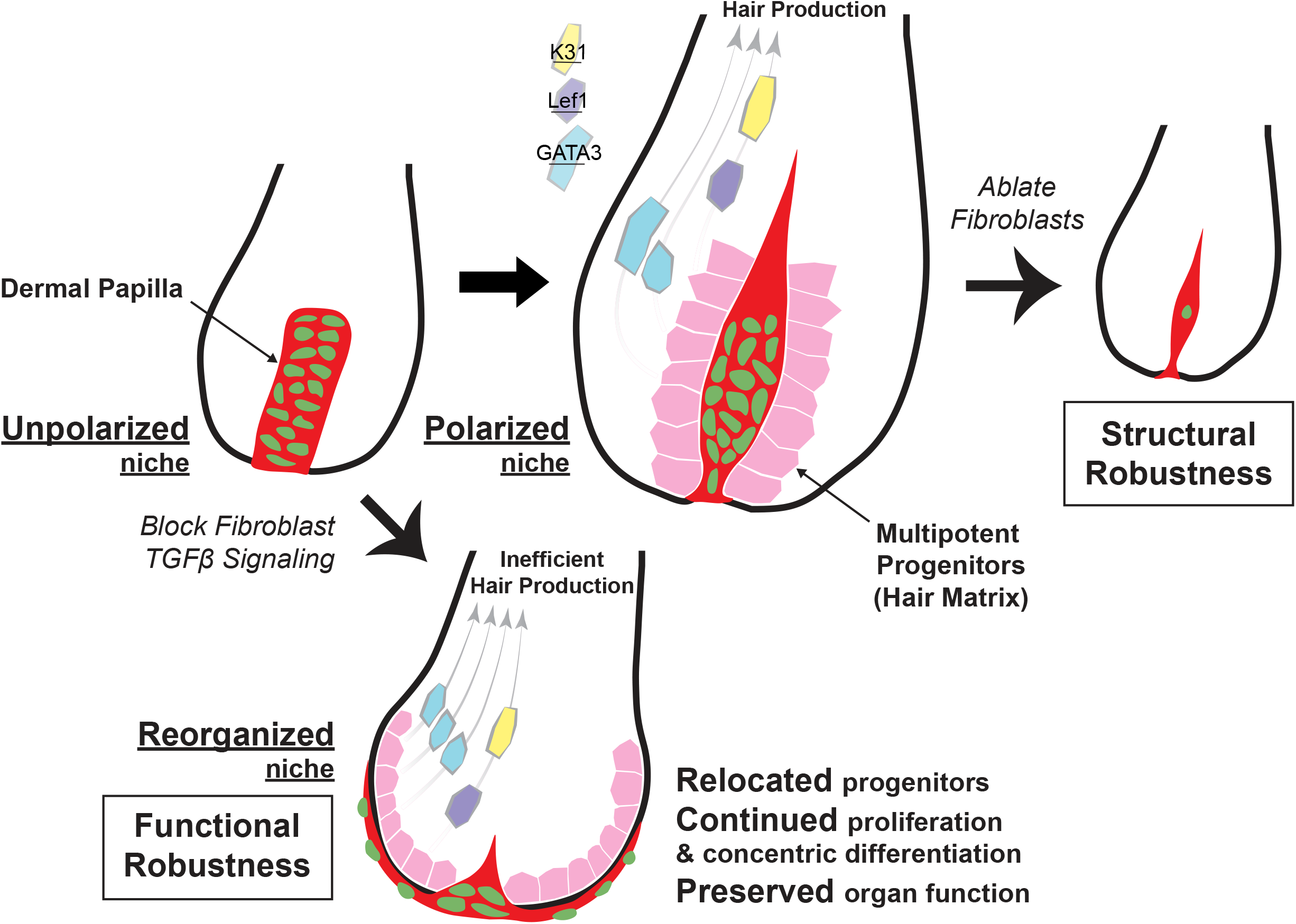
Reorganizing niche architecture still preserves organ function in the hair follicle. During hair follicle growth, the dermal papilla niche remodels into an upward polarized architecture that confers structural robustness even in the face of fibroblast death. The architecture and location of the dermal papilla are actively maintained through fibroblast TGFβ signaling. In a reorganized sheath-like niche architecture, multipotent progenitors relocate to the outer layers of the epithelium but the niche remains functionally robust to support their proliferation and concentric differentiation program. However, differentiation is completed prematurely, resulting in less efficient hair production.

Our 3D reconstruction and longitudinal analyses reveal that dermal papilla fibroblasts acquire long membrane protrusions by the late growth (polarized) stage (Fig.1), allowing them to contact several adjacent and distant hair progenitors. Recent transcriptomic studies have led to a model whereby heterogeneous fibroblasts are organized into micro-niches to instruct different differentiation trajectories of neighboring progenitors (Adam et al., 2018, Joost et al., 2020, Shin et al., 2020). Fibroblasts with extended membrane protrusions could form local micro-niche crosstalk if signaling factors are directionally transported along those protrusions, as in fly wing imaginal disc and germline stem cells (Ramirez-Weber and Kornberg, 1999, Inaba et al., 2015). Alternatively, micro-niches could be formed by extended fibroblasts if the potential of fate specification differs between large (cell body) and smaller (membrane protrusion) fibroblast-progenitor contact areas, similar to cell fate specification in ascidian embryogenesis (Guignard et al., 2020). The latter model could explain why cell bodies and cell membranes generally occupy different compartments within the polarized dermal papilla (Fig.1). Future investigation into the formation and function of niche polarity will illuminate how the niche coordinates its constituent cells to remodel different niche compartments and perform diverse niche functions.

Here, we discovered a new role for TGFβ signaling in dermal papilla fibroblasts to actively maintain niche architecture, in contrast to the common assumption that it is passively remodeled by the epithelium. Previous manipulations of dermal papilla signaling status and identity signatures altered activities of multipotent progenitors, but did not noticeably affect the architecture of the niche (Enshell-Seijffers et al., 2010, Clavel et al., 2012, Chi et al., 2017, Harshuk-Shabso et al., 2020). In contrast, we show that genetically deleting TGFβ receptor 1 in dermal papilla fibroblasts affects multipotent progenitor activity by reorganizing niche fibroblasts which continue to express most niche markers, such as Sox2 and Integrin alpha9 (Fig.2). However, we noted that Lef1 is down-regulated in the reorganized dermal papilla (Fig.3D). Together, these results imply that niche identity is at least partially separable from its architecture. Furthermore, TGFβ signaling becomes activated just prior to the niche’s structural polarization, indicating that distinct regulatory mechanisms govern each step of niche architecture remodeling. Discovering additional regulators of this remodeling process will allow us to disentangle the functional relevance of multiple aspects of niche architecture and progenitor organization.

Finally, organ architecture and function are thought of as intimately connected, so it is surprising that dermal papillae which are re-organized into a sheath are nevertheless able to support hair progenitor proliferation and differentiation (Fig.3). Even more strikingly, reorganized dermal papillae support progenitor differentiation in a concentric organization resembling wild-type hair follicles and mirroring the concentric organization of prospective epithelial stem cells during hair follicle morphogenesis (Morita et al., 2021). This surprising result suggests that at the late growth stage, niche architecture maintenance reinforces and optimizes pre-established progenitor differentiation programs. Alternatively, this result may suggest that the purpose of dermal papilla enclosure and polarized elongation at mid-to late-growth stages is primarily to maintain proximity to differentiating multipotent progenitors, and a sheath-like niche would only be proximal to progenitors at the beginning of their differentiation journey. These models prompt new frameworks in which to scrutinize the functional importance of niche architectures in different organs (e.g., sheath-like niches in sweat glands, lung, and intestines (Shyer et al., 2015, Lu et al., 2016, Shoshkes-Carmel et al., 2018, Rao-Bhatia et al., 2020, Cordero-Espinoza et al., 2021, Goodwin et al., 2022)) and across species (e.g., sheath-like niches in the *Drosophila* germline, enclosed niches in feather follicles and human epidermal rete ridges (Losick et al., 2011, Solanas and Benitah, 2013, Wu et al., 2021)). Learning the functional principles of niche architecture from these diverse biological systems will not only deepen our understanding of when and how organ architecture affects its function, but will also provide insights about designing cell therapies and engineered tissues with efficiently coupled architecture and function.

### Limitations of study

In this study, we focused on the entire dermal papilla architecture of the *Tgfbr1*^*fl/fl*^ mutant in its growth stage at the first hair cycle. Future work will study the effects of *Tgfbr1* removal on the morphology of individual dermal papilla fibroblasts. Furthermore, to understand how niche architecture loss affects hair follicle function long-term, it will be important to examine the impact of reorganized niche architecture during hair follicle regression and the next hair cycle. Finally, *Pdgfrα*-CreER is a pan-fibroblast driver, which limits our ability to understand the contribution of other fibroblast populations to dermal papilla architecture.

## Materials and Methods

### Mice

*Pdgfrα*-H2BGFP (JAX Strain #007669) (Hamilton et al., 2003), *Pdgfrα*-CreER (JAX Strain #018280) (Kang et al., 2010), Rosa26-loxP-membraneTomato-(stop)-membraneGFP (mTmG) (JAX Strain #007676) (Muzumdar et al., 2007), ROSA-LSL-DTA (JAX Strain #009669) (Voehringer et al., 2008), R26-LSL-tdTomato mice (JAX Strain #007909) (Madisen et al., 2010) were obtained from The Jackson Laboratory. *Tgfbr1*^*fl/fl*^ mice were obtained from V. Kaartinen (Larsson et al., 2001).

mNeonGreen-Smad4 (mNG-Smad4) mice were generated as follows. CRISPR-Cas9 guide RNA was targeted to a site 43bp downstream from the endogenous Smad4 translation start site in exon 2. The linear DNA gBlock repair template (Integrated DNA Technologies) consisted of mNeonGreen (708bp), a linker (12bp), a CRISPR target site recoded with silent mutations to prevent recombination of the edited sequence, flanked by 36bp homology arms. Following PCR amplification and purification of the gBlocks, DNA, CRISPR-Cas9 nuclease, and guide RNAs were introduced by pronuclear injection into B6SJL embryos then transferred to pseudo-pregnant recipient CD1 females (JHUSOM Transgenic Core). Pups carrying the integrated mNG-Smad4 sequence were identified by genomic PCR amplification of the insertion site. As homozygote progeny could not be produced, founder lines were bred into a CD1 background (JAX) and maintained as heterozygotes. (Sequence information in SI.2)

For validation experiments in primary mouse embryonic fibroblasts (MEFs), MEFs were derived from heterozygous mNG-Smad4 mice and wild-type littermates using previously described protocols (Pokrass et al., 2020). Excluding the brain, liver, and heart, tissue from dissected E13.5 embryos was dissociated by overnight incubation in 0.25% trypsin-EDTA and plated in MEF culture medium (DMEM with 15% fetal bovine serum (FBS), 100U/ml penicillin and 100U/ml streptomycin). For imaging experiments, 7×10^3^ MEFs were seeded to fibronectin (EMD-Millipore)-coated 96-well glass-bottom plates (CellVis). Cells were imaged at 5-min intervals; vehicle control or TGF-β1 (R&D #7754-BH, to an end concentration of 10ng/μl) was added to media after 20 minutes of imaging onset. Live specimens were maintained at 37°C in 5% CO_2_ in a humidified environmental control chamber (OKOlab) during imaging. Images were acquired on a Metamorph-controlled Nikon Eclipse Ti epifluorescence microscope fitted with a 40X oil objective and a Photometric Prime 95B CMOS camera.

To label fibroblasts in a mosaic pattern, we benefited from the endogenous leakage of *Pdgfrα*-CreER; mTmG mice within the fibroblast population. Therefore, even without tamoxifen induction, a small population of fibroblasts was labeled and we could define their morphology. To achieve diphtheria toxin-induced fibroblast ablation, five doses of 100mg/kg tamoxifen in corn oil were delivered via intraperitoneal injection into *Pdgfrα*-H2BGFP; *Pdgfrα*-CreER; mTmG; ROSA-LSL-DTA mice. Doses were given twice per day for three days starting at postnatal day 24-25 (including both mid and late-growth hair follicles). To achieve conditional knockout of Tgfbr1 allele in fibroblasts, three doses of 100mg/kg tamoxifen in corn oil were delivered via intraperitoneal injection into *Pdgfrα*-CreER; *Tgfbr1*^*+/fl*^ and *Tgfbr1*^*fl/fl*^ mice. Doses were given once per day for three days, starting at postnatal day 18-20 (quiescent telogen hair follicles).

Mice of either sex were randomly selected for experiments. No blinding was done. All studies and procedures involving animal subjects were approved by the Institutional Animal Care and Use Committee at Yale School of Medicine and we conducted in accordance with the approved animal handling protocol no. 11303. This study is compliant with all relevant ethical regulations regarding animal research.

### Intravital imaging and tracking hair growth

Procedures of intravital imaging were similar to those previously described (Pineda et al., 2015). During the experiment, we maintained mice anaesthetized via vaporized 1.5% isoflurane and 1.5% oxygen in air and placed them on a warming pad to maintain body temperature. The mouse ear was mounted on a speculum with a glass coverslip on top. We used a LaVision TriM Scope II (LaVision Biotec) microscope equipped with a Chameleon Vision II (Coherent) two-photon laser (940nm) and a Chameleon Discovery (Coherent) two-photon laser (1120nm) for intravital imaging. Either a Zeiss ×20/1.0, Nikon ×25/1.10 or a Nikon ×40/1.15 water immersion objective was used to scan a field of view of 500μm^2^ (20X), 505μm^2^ (25X), or 278μm^2^ (40X). Via a motorized stage, we acquired multiple tiles of z-stacks that reach up to 200μm deep and vary each z step from 1 to 10μm depending on the requirement of spatial resolution. Patterns of hair follicles were used as landmarks to revisit the same skin area longitudinally.

Timing of hair cycle initiation varies slightly between different mice. If necessary, we kept imaging until the mice reached the stage of interest. Wild-type and *Tgfbr1*^*+/fl*^ control hair follicle stages were identified based on previous literature (MuÈller-RoÈver et al., 2001). In general, early-growth stages include anagen 1 to anagen 2 (early); mid-growth stages include anagen 2 (late) to anagen 3b; late-growth stages include anagen 3c-6. *Tgfbr1*^*fl/fl*^ mutant hair follicle stages were identified based on dermal papilla morphology. Mutant dermal papillae at Stage1 look similar to anagen 3a-b in control littermates. By contrast, mutant dermal papillae at Stage2 are triangle-shaped, at Stage3 are very thin, and disappear from within the epithelium at Stage4. The frequency and length of longitudinal revisits depended on the purpose of each experiment and is indicated in the figures and legends.

To track hair growth, we shaved the back skin and removed hair using Nair Hair Removal Cream once during postnatal day 18-21. Then, these mice were photographed on the next day and every week until the end of first hair cycle at postnatal day 60-63. During all procedures, mice were anaesthetized and kept on a warming pad similar to intravital imaging.

### Immunofluorescence

Based on hair follicle stages of interests, *Tgfbr1*^*+/fl*^ control back skin samples were collected at postnatal day 34-35, and *Tgfbr1*^*fl/fl*^ mutant back skin samples were collected at postnatal day 37. For thick tissue sections, back skins were dissected, fixed in 4% paraformaldehyde (PFA) in PBS at room temperature for an hour and then 30% sucrose at 4°C overnight. These samples were embedded in optimal cutting temperature compound (OCT) and frozen on dry ice. The OCT blocks were later sectioned at 30μm thickness to preserve 3D architecture. These fixed sections were further fixed in 4% PFA for 20 minutes and washed in 2% Triton X-100 in PBS three times (each 10 minutes) at room temperature. Then the sections were incubated in the blocking solution (5% normal donkey serum, 1% BSA and 2% Triton X-100 in PBS) at room temperature overnight. Later, the sections were incubated with primary antibodies diluted in the blocking solution at room temperature for one day, and then washed in 2% Triton X-100 in PBS three times (each 10 minutes) at room temperature. The sections were further incubated with secondary antibody diluted in the blocking solution at room temperature overnight and then washed in 2% Triton X-100 in PBS three times (each 10 minutes). The stained sections were mounted in SlowFade Diamond Antifade Mountant (S36972, Invitrogen) and imaged under the two-photon microscope.

The following primary antibodies were used: human/mouse active Caspase3 (AF835, R&D Systems, 1:200), mouse integrin alpha9 (AF3827, R&D Systems, 1:200), Sox2 monoclonal antibody, eBioscience (14-9811-82, Invitrogen, 1:400), Lef1 monoclonal antibody (E.281.8) (MA5-14966, Invitrogen, 1:400), recombinant anti-GATA3 antibody [EPR16651] (ab199428, Abcam, 1:200), anti-acidic hair keratin K31 guinea pig polyclonal (GP-hHa1, Progen, 1:200). The following secondary antibodies were used: goat anti-rabbit Alexa 488 (A32731, Invitrogen, 1:200), donkey anti-goat Alexa 488 (A-11055, Invitrogen, 1:200), donkey anti-rat Alexa 488 (A-21208, Invitrogen, 1:200), goat anti-guinea pig Alexa 488 (A-11073, Invitrogen, 1:200).

*Tgfbr1*^*+/fl*^ and *Tgfbr1*^*fl/fl*^ ear skin was collected at postnatal day 39. For whole-mount staining, ear skin was dissected and fixed in 4% PFA in PBS at 4°C overnight. Fixed tissues were cut into small pieces and incubated in the blocking solution (5% normal donkey serum, 1% BSA and 2% Triton X-100 in PBS) at 37°C for 3 days. Blocked tissues were incubated with anti-Ki67 antibody (ab15580, Abcam, 1:200) at 37°C for 6 days. Tissues were then washed in PBS with 2% Triton X-100 at 37°C for 4 days and incubated with goat anti-rabbit Alexa 633 (A-21071, Invitrogen, 1:100) at 37°C for 5 days. After staining, tissues were washed in PBS with 2% Triton X-100 at 37°C for 5 days and mounted in SlowFade Diamond Antifade Mountant (S36972, Invitrogen). The stained whole mounts were imaged under the two-photon microscope.

### Image analysis

Z-stacks of images were imported into Fiji (ImageJ, NIH) or Imaris (BitPlane) for analysis. Tiles of different z-stacks were stitched in Fiji. Either selected single optical z slice or summed z slices were used in figures, as indicated in legends. Quantification of percentages, measurement of length and width were performed in Fiji based on z-stack’s 3D information. 3D reconstructed dermal papillae were performed in Imaris via its surface rendering function.

### Statistics and reproducibility

Statistical calculations were performed with Prism 9 (GraphPad). P value <0.05 is considered significant; the precise P values and specific statistical tests can be found in figures and legends. Sample sizes are indicated in figure legends as well. No statistical method was used to predetermine sample size. All experiments were repeated at least three times.

## Supporting information

Supplemental Item 1

## Acknowledgements

We thank the Greco lab members, thesis committee members of H.W. (Stefania Nicoli, Haifan Lin, and Scott Holley), as well as Maria Jose Ortuno Romero, Emily Forrest, Andrew Lee Cox, and Caroline Hendry for helpful discussions and thoughtful feedbacks on the manuscript. H.W. is supported by China Scholarship Council (CSC) Grant. This work is supported by an HHMI Scholar award (55108527) and NIH grants number R01AR063663, R01AR072668, DP1AG066590 (all to V.G.)

## Author contributions

H.W., T.X. and V.G. designed the experiments. H.W. performed experiments and analyzed the data. H.W., T.X., L.E.G. and V.G. wrote the manuscript. S.D. provided help to maintain the mouse colony. J.P. and S.R. generated the mNeonGreen-Smad4 mouse line. H.A.P. helped with mutant phenotype analyses.

---

**Supplemental Figure 1.**
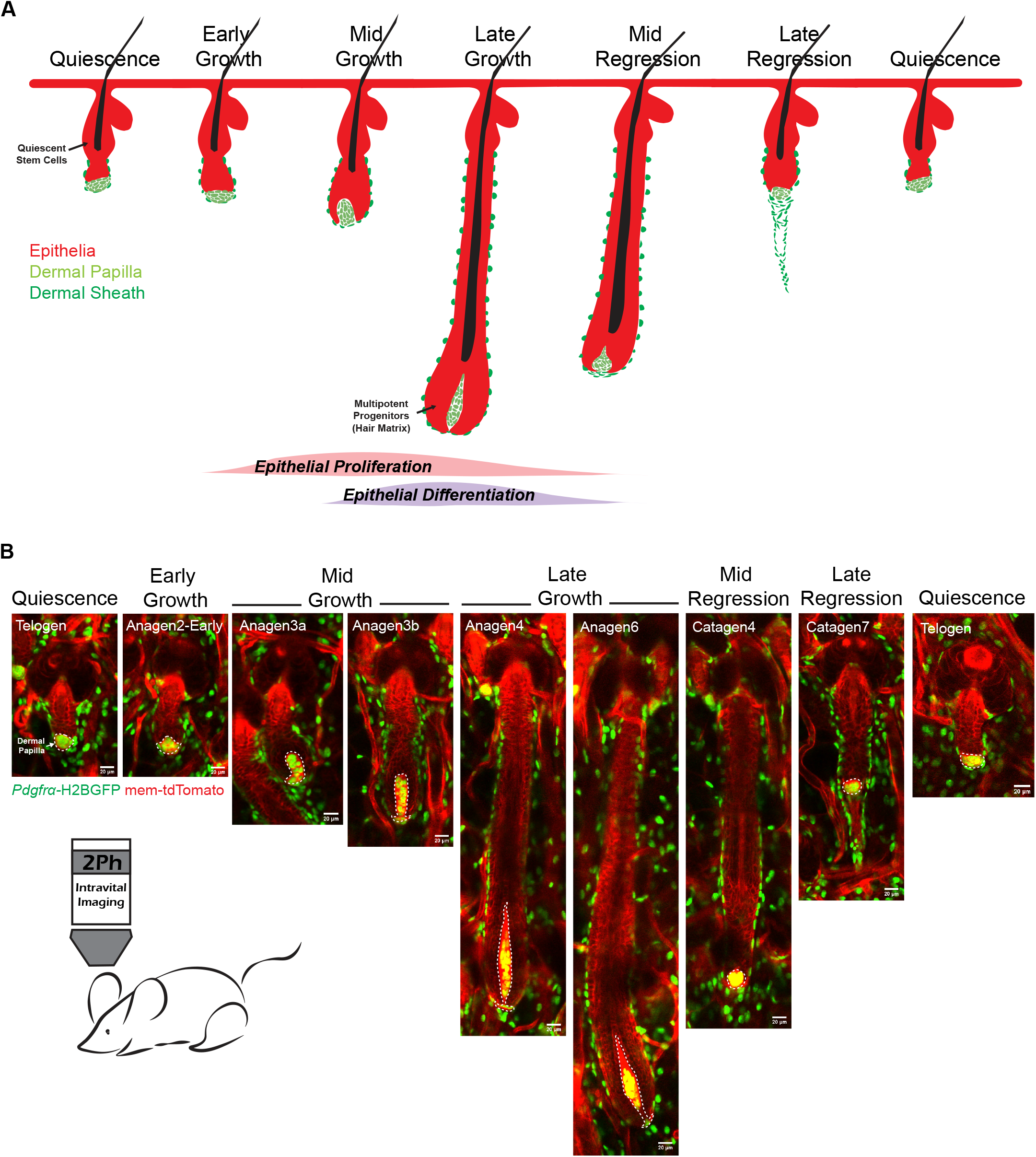
Intravital imaging of hair follicle epithelium and associated fibroblasts over an entire hair cycle, related to Figure 1. A. Schematic of the remodeling hair follicle epithelium (red) and associated fibroblasts, including dermal sheath (dark green) and dermal papilla (light green). Time windows of epithelial proliferation and differentiation are indicated in red and purple. B. Intravital images of hair follicles at different stages. *Pdgfrα*-H2BGFP labels all fibroblast nuclei in green and membrane-tdTomato labels all cell membranes in red. The dermal papillae are white dash-lined at the bottom of hair follicles. As the hair follicle grows, the dermal papilla becomes enclosed, elongated, and returns to a compact ball architecture during regression. All representative images are single optical z slice with scale bar sizes indicated.

**Supplemental Figure 2.**
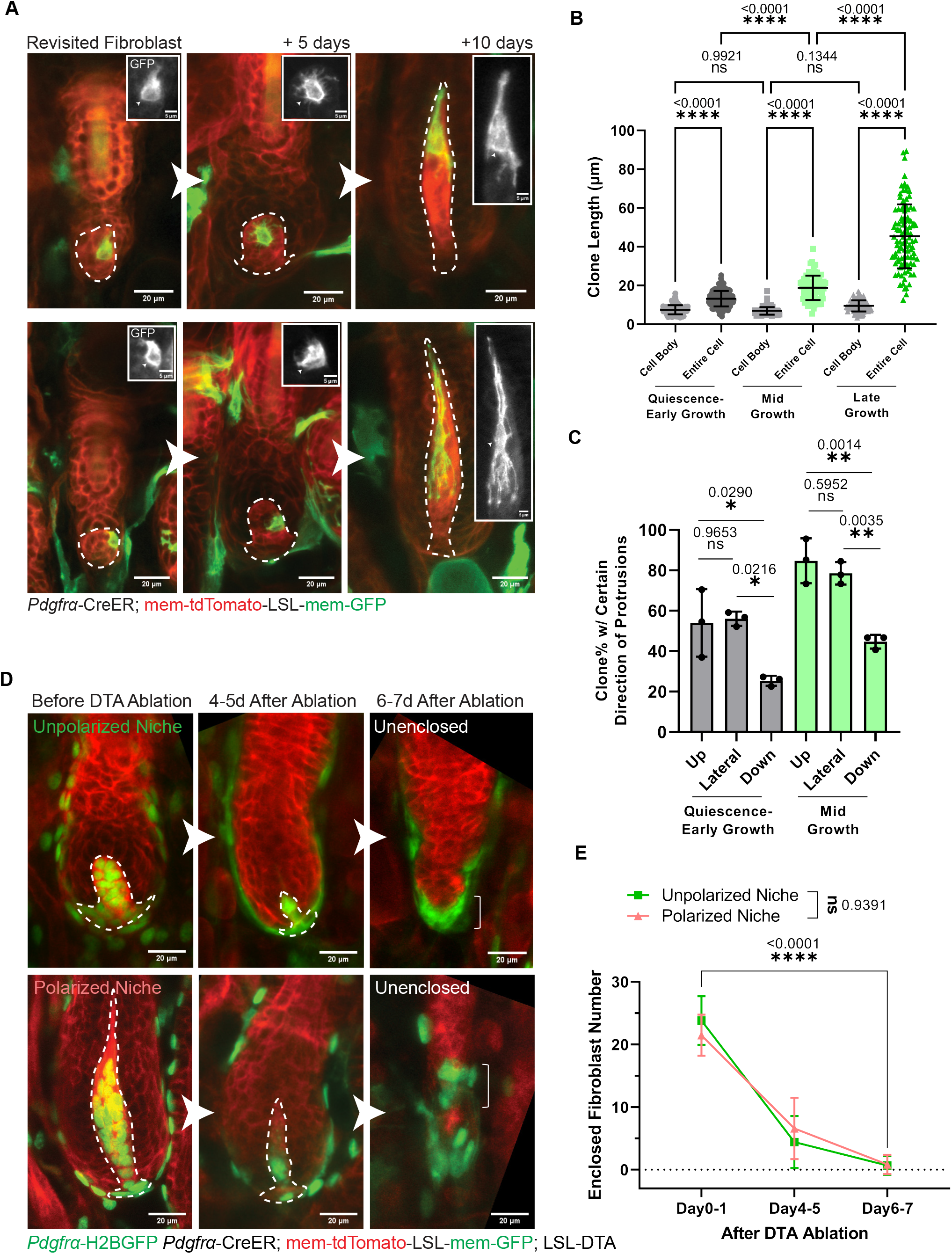
Remodeling of dermal papilla fibroblasts during hair follicle growth, and examples of unenclosed dermal papillae after fibroblast ablation, related to Figure 1. A. The same dermal papilla fibroblast longitudinally imaged from quiescence to late-growth. A single fibroblast is labeled in green by *Pdgfrα*-CreER; mTmG within the dash-lined dermal papilla. Insets are crops of fibroblasts with membrane-GFP in gray and cell body indicated with arrowheads. Representative images are summed optical z slices with scale bar sizes indicated. B. Length of dermal papilla fibroblast cell body and entire cell membranes at different hair cycle stages. Data are presented as mean ± S.D. n=160 dermal papilla fibroblast clones at quiescence-early growth from 3 mice. n=88 dermal papilla fibroblast clones at mid-growth from 3 mice. n=103 dermal papilla fibroblast clones at late-growth from 3 mice. Tukey’s multiple comparisons test. C. Percentage of dermal papilla fibroblasts harboring certain direction of membrane protrusions (upward, lateral, downward) at quiescence-early growth and mid-growth. Data are presented as mean ± S.D. n=161 dermal papilla fibroblast clones at quiescence-early growth from 3 mice. n=87 dermal papilla fibroblast clones at mid-growth from 3 mice. Tukey’s multiple comparisons test. D. Longitudinally imaged dermal papillae that become unenclosed after diphtheria toxin (DTA)-induced fibroblast ablation (*Pdgfrα*-CreER; LSL-DTA). Fibroblast nuclei are labeled in green by *Pdgfrα*-H2BGFP, and cell membranes are in red by membrane-tdTomato. Before DTA ablation and 4-5 days after ablation, dash-lined dermal papillae are enclosed within the epithelium; some of these become unenclosed and locate at the bottom of hair follicle epithelium 6-7 days after ablation. Representative images are summed optical z slices with scale bar sizes indicated. E. The number of fibroblasts that remain enclosed within the hair follicle epithelium after DTA-induced fibroblast ablation. Fibroblast number is counted based on green fibroblast nuclear signal labeled by *Pdgfrα*-H2BGFP. Data are represented as mean ± S.D. For unpolarized dermal papillae at mid-growth, n=41 (Day0-1), 53 (Day4-5), and 60 (Day6-7) from 3 mice; for polarized dermal papillae at late-growth, n=72 (Day0-1), 78 (Day4-5), and 71 (Day6-7) from 3 mice. Mixed-effects analysis.

**Supplemental Figure 3.**
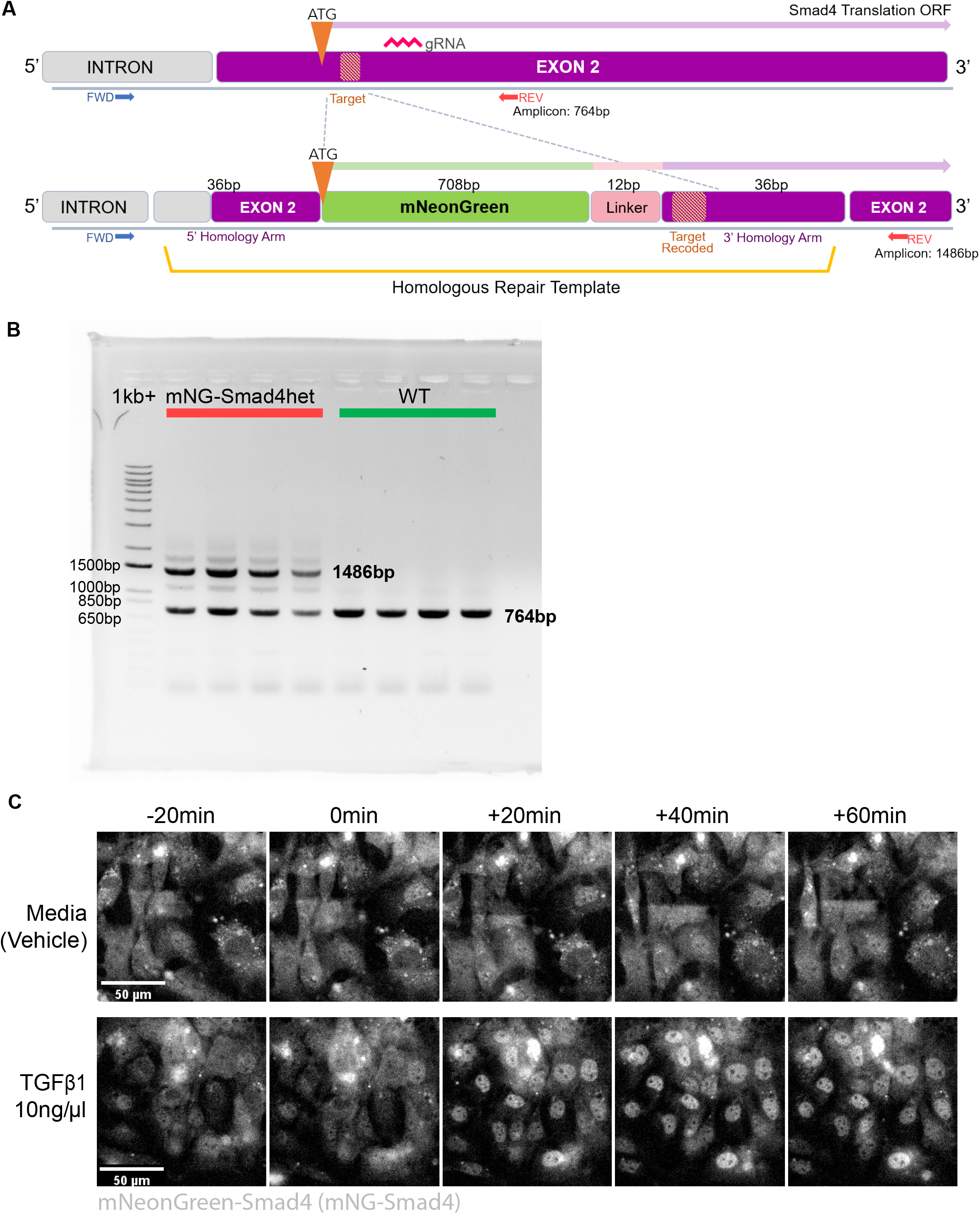
Reporter mouse generation by endogenous N-terminal tagging of Smad4, related to Figure 2. A. Schematic of the endogenous Smad4 locus in wild-type (above) and mNeonGreen-Smad4 mouse (below) following CRISPR-Cas9-guided homology-directed repair. mNeonGreen (mNG) and a 12bp linker sequence were inserted immediately upstream of the Smad4 start codon in exon 2, flanked by 36bp homology arms. Codons in the CRISPR gRNA target and PAM sequence were replaced with silent mutations to prevent further recombination. Red and blue arrows indicate position of genotyping PCR oligos and expected amplicon size. B. Representative gel electrophoresis image showing genotyping PCR of mNG-Smad4. In addition to the wild-type 764bp amplicon (green), heterozygous mNG-Smad4 mice have an additional 1486bp band (red) indicating the integrated repair template. C. Representative still images of mouse embryonic fibroblasts derived from mNG-Smad4 heterozygote mice. 20 minutes into imaging, live cells were stimulated with TGF-β1 (10ng/μl) or vehicle. Images were acquired every 5min and are shown here at 20min intervals. Nuclear localization of mNG-Smad4 remained stable for approximately 7 hours following stimulation.

**Supplemental Figure 4.**
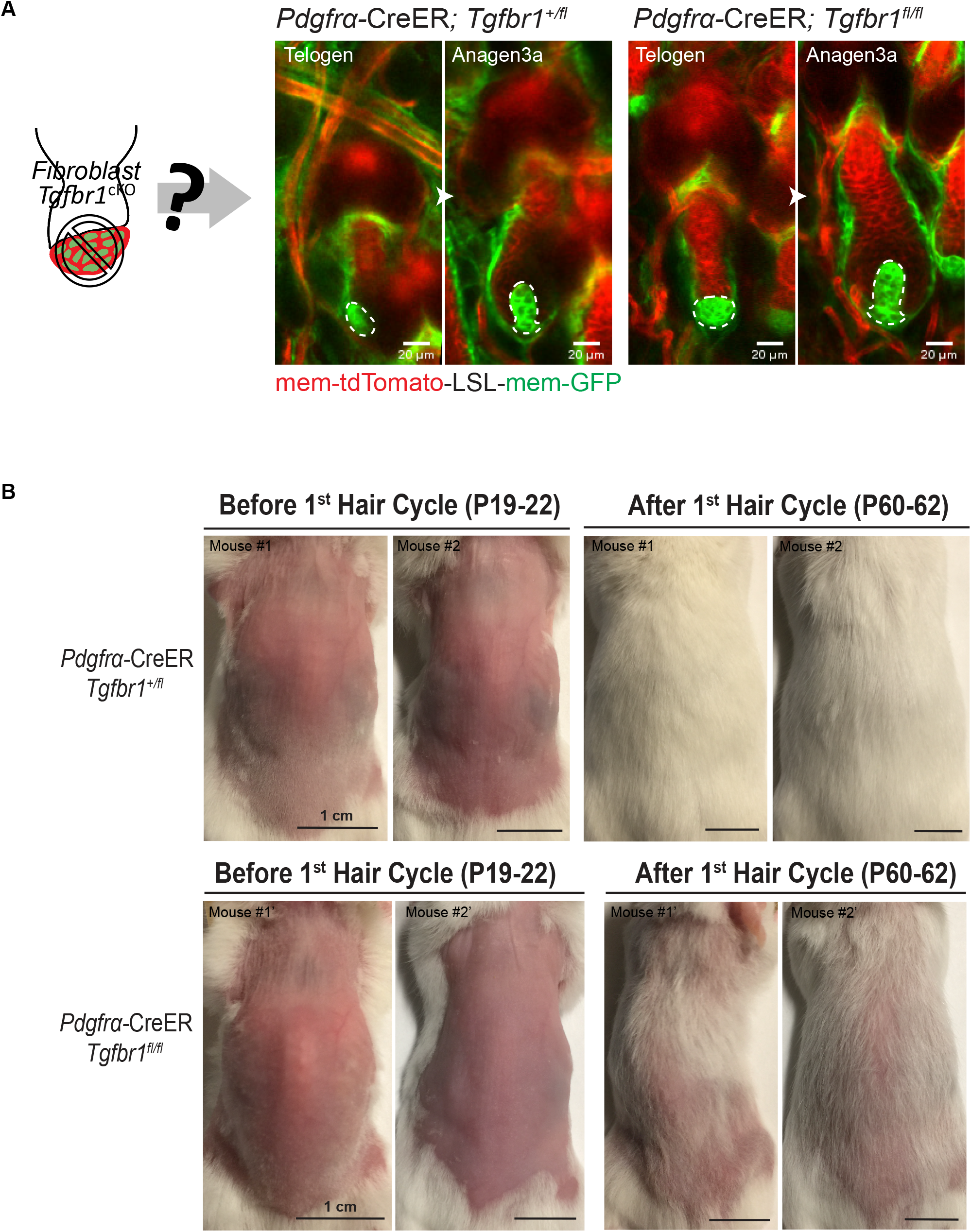
Hair follicles with *Tgfbr1*^*fl/fl*^ mutant fibroblasts proceed to mid-growth but produce hairs less efficiently, related to Figure 2-3. A. Revisits at quiescence (Telogen) and mid-growth (Anagen 3a) of the same *Tgfbr1*^*+/fl*^ or *Tgfbr1*^*+/fl*^ dermal papillae, after *Pdgfrα*-CreER induction in quiescence. The dermal papillae remain enclosed within the epithelium until mid-growth. Fibroblast membranes are labeled in green membrane-GFP and other membranes are in red membrane-tdTomato. Dermal papillae are dash-lined. Representative images are single optical z slice with scale bar sizes indicated. B. Representative photos of additional replicate mice (*Tgfbr1*^*+/fl*^ and *Tgfbr1*^*fl/fl*^) with back skin shaved at the quiescent stage, and revisited after an entire first hair cycle. Scale bars, 1 cm.

**Supplemental Item 1: Movie of a three-dimensionally reconstructed dermal papilla and individual fibroblasts, related to Figure 1**

The entire dermal papilla at late-growth is 3D reconstructed based on its membrane-tdTomato signal. Within the red outlined dermal papilla, cell bodies of individual fibroblasts are rendered into surfaces with different colors. Further, a single fibroblast is 3D reconstructed based on its membrane-GFP signal. The cell body of this fibroblast is rendered into a pink surface and its entire membrane in a green surface.

**Supplemental Item 2: Sequence information for mNeonGreen-Smad4 mice**

## gRNA for CRISPR

ACTATGTACAATGCTCAGAC (20bp, REV)

### HDR template (837bp linear DNA gBlock, from Integrated DNA Technologies)

*Capitalized sequences at either end denote the 36bp 5’ & 3’ homology arms. Lowercase denotes the inserted mNeonGreen and linker sequences. Capitalized bases preceding the 3’ homology arm denote silent mutations in the CRISPR-Cas9 target area, specifically residues downstream of the original SMAD4 start codon up to the gRNA PAM site*.

TTGGAGAGTTGGATTTAAAAGAAAAAACTTGAACAAatggtgagcaagggcgaggaggataacatggcctctctcccag

cgacacatgagttacacatctttggctccatcaacggtgtggactttgacatggtgggtcagggcaccggcaatccaaatgatggttatgaggagtt

aaacctgaagtccaccaagggtgacctccagttctccccctggattctggtccctcatatcgggtatggcttccatcagtacctgccctaccctgacg

ggatgtcgcctttccaggccgccatggtagatggctccggctaccaagtccatcgcacaatgcagtttgaagatggtgcctcccttactgttaactac

cgctacacctacgagggaagccacatcaaaggagaggcccaggtgaaggggactggtttccctgctgacggtcctgtgatgaccaactcgctg

accgctgcggactggtgcaggtcgaagaagacttaccccaacgacaaaaccatcatcagtacctttaagtggagttacaccactggaaatggc

aagcgctaccggagcactgcgcggaccacctacacctttgccaagccaatggcggctaactatctgaagaaccagccgatgtacgtgttccgta

agacggagctcaagcactccaagaccgagctcaacttcaaggagtggcaaaaggcctttaccgatgtgatgggcatggacgagctgtacaag

tccggatccagaatggaTaaCatgtcGatTacCaaCacCccGacCagCaaTgaCgcATGTCTGAGCATTGTACATAGT

TTGATGTGTCATAGA

### PCR primers

DNA oligos (ordered from Integrated DNA Technologies, stored as 200μM stock)

FWD: GCC TTT GCA ATA GTG AGG AGA CCG G

REV: GCA TCT GGG AAT GCT CTC TTC TCG CC

